# LSD induces increased signalling entropy in rats’ prefrontal cortex

**DOI:** 10.1101/2021.06.23.449556

**Authors:** Aurora Savino, Charles D. Nichols

## Abstract

Psychedelic drugs are gaining attention from the scientific community as potential new compounds for the treatment of psychiatric diseases such as mood and substance use disorders. The 5-HT_2A_ receptor has been identified as the main molecular target, and early studies pointed to an effect on the expression of neuroplasticity genes. Analysing RNA-seq data from the prefrontal cortex of rats chronically treated with lysergic acid diethylamide (LSD), we describe the psychedelic-induced rewiring of gene co-expression networks, which become less centralized but more complex, with an overall increase in signalling entropy, typical of highly plastic systems. Intriguingly, signalling entropy mirrors, at the molecular level, the increased brain entropy reported through neuroimaging studies in human, suggesting the underlying mechanisms of higher-order phenomena. Moreover, from the analysis of network topology we identify potential transcriptional regulators and imply different cell types in psychedelics’ activity.

## 1. Introduction

Psychedelics, such as lysergic acid diethylamide (LSD), psilocybin and the substituted amphetamine 1-(2,5-dimethoxy-4-iodophenyl)-2-aminopropane (DOI), are psychoactive compounds that induce profound acute subjective effects in humans, affecting perception, behaviour, and mood through activation of 5-HT_2A_ receptors (D. E. Nichols, 2004). The alterations in perception and thought processes can be interpreted as transient psychotic-like states (D. E. Nichols, 2016) that resemble symptoms of mental disorders such as schizophrenia (Schmid et al., 2015; F X Vollenweider et al., 1998). Hence, they have been used to mimic schizophrenia in animal models (Marona-Lewicka et al., 2011; Martin et al., 2014), similarly to the previously employed NMDA receptor antagonists ketamine and PCP (Winship et al., 2019). However, psychedelics have also been recently shown to produce long-lasting beneficial effects in patients with mood and substance use disorders: several pilot clinical trials have shown that single acute administrations of psilocybin or LSD produce symptoms’ reduction for at least six months (R. L. Carhart-Harris et al., 2018; Robin L. Carhart-Harris et al., 2016, 2017; Davis et al., 2020; Griffiths et al., 2016; Santos et al., 2016), contrary to most classical therapeutics for mood disorders (e.g. serotonin reuptake inhibitors), which require weeks to exert their effects. Moreover, single administrations of psilocybin produce long-lasting changes in personality traits (e.g. higher *openness*) that extend also to healthy subjects (Erritzoe et al., 2018; MacLean et al., 2011). Importantly, these treatments have been proven safe, without subsequent persisting psychosis, drug abuse or any significant impairment in cognitive or physical functioning (Studerus et al., 2011). Nevertheless, in rare instances perceptual alterations can reoccur as “benign flashbacks” (Martinotti et al., 2018), also known as hallucinogen persisting perception disorder (HPPD).

The molecular target responsible for psychedelics’ behavioural and psychological effects in humans has been identified as the 5-HT_2A_ receptor (Madsen et al., 2019; Preller et al., 2018; Franz X. Vollenweider & Preller, 2020). Nevertheless, molecular mechanisms underlying acute changes in brain activity and lasting psychological effects have been less investigated. Neuroplasticity-related genes and immediate early genes have been reported to be over-expressed upon psychedelics’ administration in rodent models, especially in areas with dense 5-HT_2A_ receptor expression like the prefrontal cortex (Benekareddy et al., 2013; González-Maeso et al., 2003, 2007; Jefsen et al., 2020; C. D. Nichols et al., 2003). These results have been confirmed and expanded through the use of high-throughput technologies, profiling the whole transcriptome, such as microarrays (C. D. Nichols & Sanders-Bush, 2002, 2004) and, more recently, RNA sequencing (Donovan et al., 2021; Martin et al., 2014; Revenga et al., 2021).

Given their relevance for basic research and the study of schizophrenia, on one hand, and as promising therapeutic agents for psychiatric disorders on the other, much research effort has been devoted to investigate their mechanisms of action. In particular, most research interest has been focussed on neuroimaging (e.g. MEG or fMRI) of subjects during psychedelics’ acute effects, leading to the observation of increased variability of spontaneous brain activity and changes in brain networks’ connectivity (Preller et al., 2018, 2019, 2020). Using these measurements, the entropy (variability or information content) of brain activity has been quantified and shown to increase upon psychedelics (Robin L. Carhart-Harris et al., 2014). Interestingly, acute increase of brain entropy under LSD correlates with changes in personality traits after two weeks (Lebedev et al., 2016). From its first formulation, this theory has gained additional support through new measurements (Robin L. Carhart-Harris, 2018; Herzog et al., 2020; Lebedev et al., 2016; Schartner et al., 2017).

More generally, entropy is a measure of the number of possible states of a system, and can manifest at different levels in signals of different kinds, not only from brain’s electrical activity but also from gene expression. In cell biology, it intuitively quantifies cell plasticity and robustness to perturbations. Indeed, stem cells have been shown to have high transcriptional entropy that decreases across differentiation (Gulati et al., 2020; Teschendorff & Enver, 2017), and tumors, in particular those resistant to drug treatments, display higher entropy than normal tissue (Conforte et al., 2019; Nijman, 2020; Savino et al., 2020). Both systems are highly plastic, able to adapt to the environment and diversify in response to stimuli. However, diversity can increase also with the loss of stable organizing principles, correlating with dysfunctions such as aging (Hernando-Herraez et al., 2019).

Here, analysing the RNA-seq data of rats chronically treated with LSD (Martin et al., 2014), we dig deeper into the impact that psychedelics exert on gene expression. We investigate how gene co-expression networks reorganize upon LSD and identify Tcf4 as a potential player in network regulation. Moreover, we find differences in network topology for microenvironmental or neuronal networks, suggesting that both cell compartments might be involved in the long-term effects of repeated LSD administration. Finally, we propose a model where the increase in brain entropy after a single dose of the drug is paralleled by increased transcriptional entropy, which declines after a few days in absence of a new intake. Repeated drug administration results in a sustained increase in transcriptional entropy, lasting at least a few weeks after discontinuation of the drug. Moreover, we identify epigenetic factors that might be responsible for these lasting effects, and the potential involvement of alternative splicing patterns and transposable elements’ activation in the overall activity of psychedelics.

## 2. Results

### 2.1. Chronic LSD exposure has a long-term effect on the epigenetic machinery

We made use of the previously described system of rats chronically treated with LSD, which show persistent altered locomotor activity and social interactions long after drug discontinuation (Marona-Lewicka et al., 2011) and have hence been proposed as a rat model system for the study of schizophrenia. RNA-seq data on the mPFC of these rats 3 weeks after cessation of drug showed persisting transcriptional effects, and an enrichment for differential regulation of schizophrenia-related genes (Martin et al., 2014).

We further pursued the investigation of these high depth RNA-seq data to study the rearrangements of gene co-expression networks in response to prolonged repeated LSD administration.

First, from the analysis of differentially expressed genes (Suppl. Table 1), we observed increased expression of genes related to neuroplasticity and neurotransmission, confirming previous reports of *bdnf* up-regulation and of increased dendritogenesis upon psychedelics treatment (Ly et al., 2018). Indeed, “dendrite development” is amongst the top significantly enriched Gene Ontology (GO) categories for up-regulated genes (Suppl. Table 2). Also, we found a significant enrichment for circadian rhythm genes, possibly underlying the alteration of sleep cycles (Barbanoj et al., 2008). Interestingly, “covalent chromatin modification” and “histone modification” showed strong enrichment for up-regulated genes, among which we found *Tet1*, involved in the erasure of DNA methylation (Wu & Zhang, 2017). This indicates that repeated psychedelics’ administration affects the epigenetic machinery, thus suggesting a mechanism for their long-term effects. Down-regulated genes were found to be mostly involved in oxidative phosphorylation (Suppl. Table 3).

We then built gene co-expression networks, to investigate potential regulatory relationships between genes and their changes upon treatment. We applied the widely used WGCNA algorithm (Weighted Gene Co-expression Network Analysis) (Zhang & Horvath, 2005) and identified eighteen clusters of genes (modules, Suppl. Table 4), six of which show differential activity between LSD and Ctrl conditions (Figure 1), as quantified through the module eigengene (ME), an ideal meta-gene representing the whole module’s expression. Modules higher in LSD-treated rats are enriched for regulation of chromatin organization, vesicle-mediated transport in synapse, and cell-cell adhesion (Suppl. Table 5), indicating that our observations on differentially expressed GO categories are robust and independent of the analysis method. Although modules down-regulated upon treatment do not show a significant enrichment for GO biological processes, they include genes with molecular functions such as GTP-binding, hydrolase activity, and structural constituents of ribosomes (Suppl. Table 6). It is important to note, however, that these are not the only enriched categories for each module (Suppl. Tables 5-6), and labels assigned based on the most significantly enriched GO categories should not be interpreted strictly to exclude other categories.

**Figure 1.**
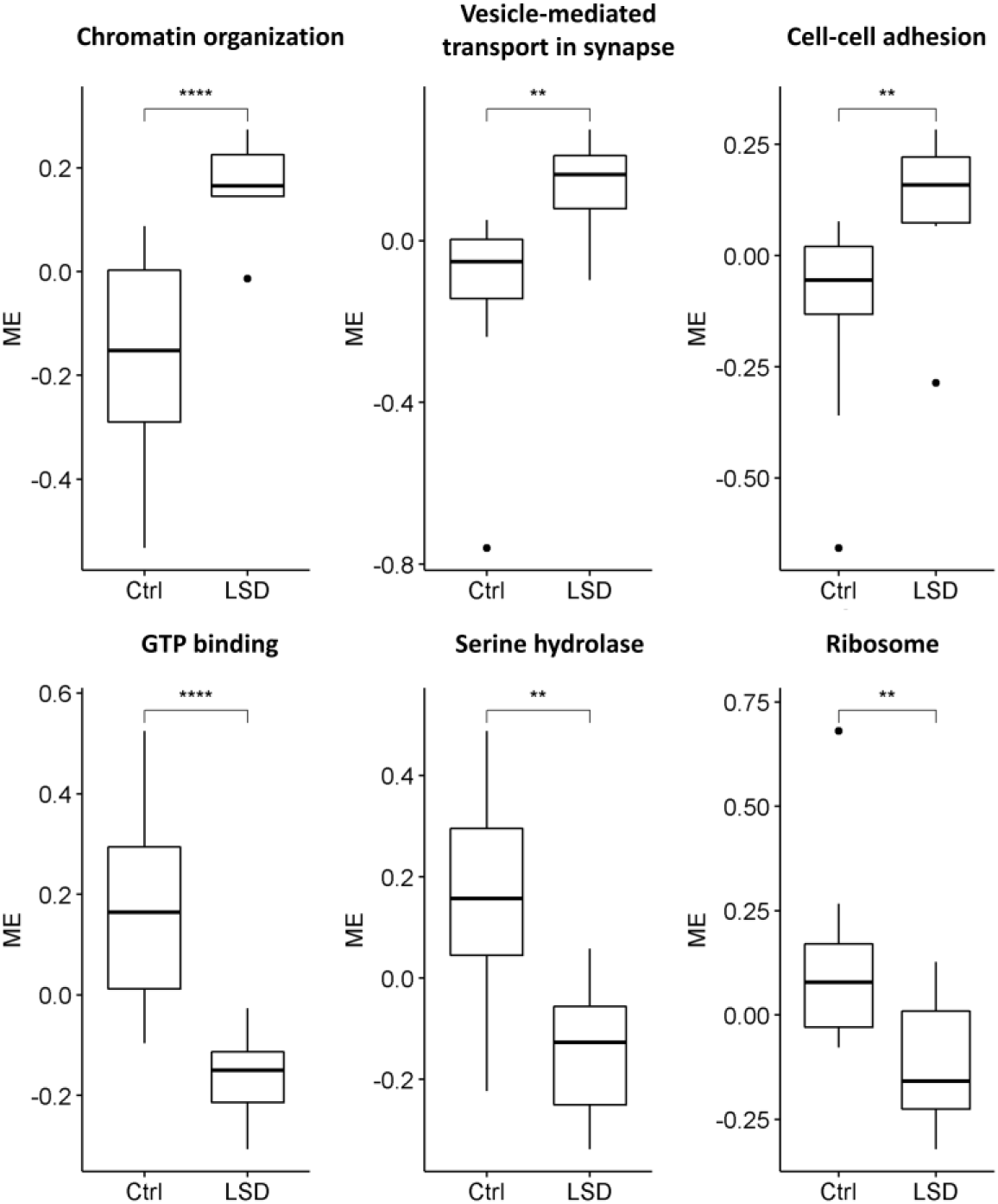
LSD-regulated co-expression modules. Three modules, enriched for the GO categories “chromatin organization”, “vesicle-mediated transport in synapse” and “cell-cell adhesion”, respectively, are more active in LSD treated rats than in controls, while the three modules enriched for the molecular functions “GTP-binding”, “serine hydrolase” and “ribosome” are less active in the LSD condition. Activity is quantified through the module eigengene (ME), the projection of samples on the first principal component obtained using the genes of the module. Significance is obtained with the Wilcoxon rank sum test. * = 0.05, ** = 0.01, *** = 0.001, **** = 0.0001

Using the network’s topological features, we looked for potential regulators of the LSD modules. Indeed, genes centrally located within the network and displaying many connections (hubs) have been reported to be crucial for the system’s maintenance (Jeong et al., 2001; Pržulj et al., 2004). Hence, we selected genes amongst the first 10 central transcription factors (TFs) in each module and tested whether there is independent evidence of their regulatory activity on the same module’s genes using the ChEA database (Lachmann et al., 2010). With these criteria, we identified Tcf4 as a potential regulator of the “Vesicle transport” module (false discovery rate for each of the three available gene sets <3*10^-4^), and Arnt as a potential regulator of the “Cell-cell adhesion” module (false discovery rate = 0.0009), but no other genes passed these stringent filters.

### 2.2. Transcriptional entropy increases with LSD treatment

We investigated the signalling variability of gene networks by measuring transcriptional entropy. As mentioned in the introduction, transcriptional entropy has been paradoxically associated with both plasticity and aging. Nevertheless, different measures have been used between the two contexts: in the first, entropy is quantified from the number of signalling paths in a protein-protein interaction network or from the number of expressed genes in each sample (Gulati et al., 2020; Teschendorff & Enver, 2017), whereas in the second, differences between samples have been used as a metric to define entropy (Hernando-Herraez et al., 2019). To distinguish these scenarios, we named these two kinds of entropies as “signalling” and “between-sample” entropy, schematically described in Figure 2.

**Figure 2.**
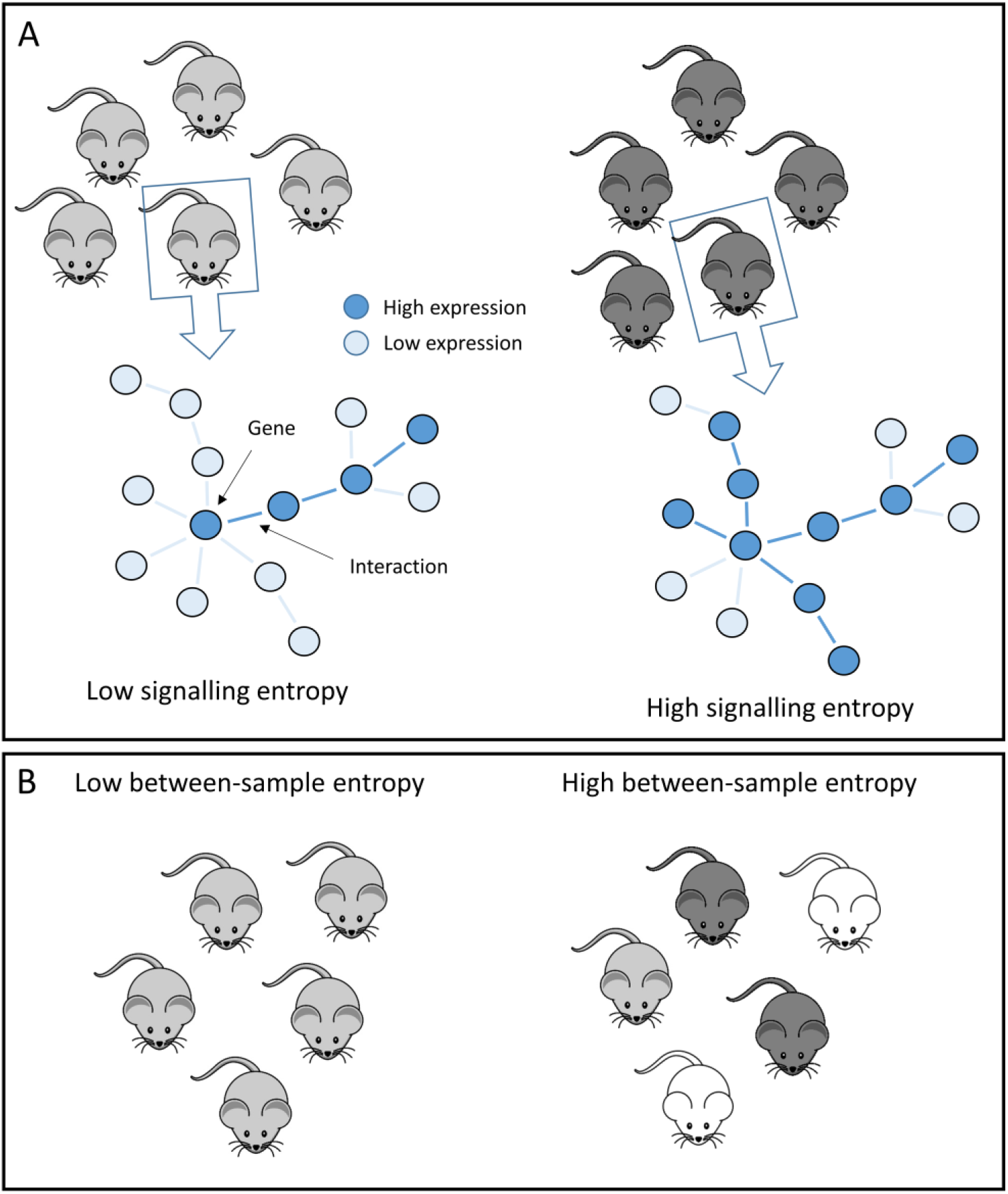
Schematic representation of signalling and between-sample entropies. A) Within each subject, a gene-interaction network is obtained and the number of possible paths from each node is calculated based on gene expression levels. A few paths correspond to low entropy, while many paths correspond to high entropy. Hence, for each subject, a gene-level entropy measure is obtained and then summarized as the overall subject specific signalling entropy. B) Between-sample entropy is defined as the variability between different subjects in a group. Hence, homogenous groups have low entropy, while diversified groups have high entropy.

To ensure robustness of our observations, we measured signalling entropy based on both protein-protein interaction (PPI) networks, as in the originally published method (Teschendorff & Enver, 2017) or on our co-expression network. Moreover, as additional measures of within-sample variability or information content, we quantified the number of splicing junctions used in each sample and the proportion of reads mapped to transposable elements. With all metrics we found a significant increase in the transcriptional complexity of the PFC of rats chronically treated with LSD with respect to saline controls, even after a withdrawal period of 3 weeks (Figure 3A-D). Of note, none of these measures are influenced by sequencing depth, since differences are retained also by randomly sampling the data to have the same number of reads in each sample (Suppl. Fig. 1). On the contrary, we observed a decrease in between-sample entropy, revealed by the lower samples’ divergence in the LSD condition at the gene expression and splicing usage levels (Figure 3E-F). Taken together, these observations suggest that LSD treatment induces increased plasticity-like and reduced aging-like entropy.

**Figure 3.**
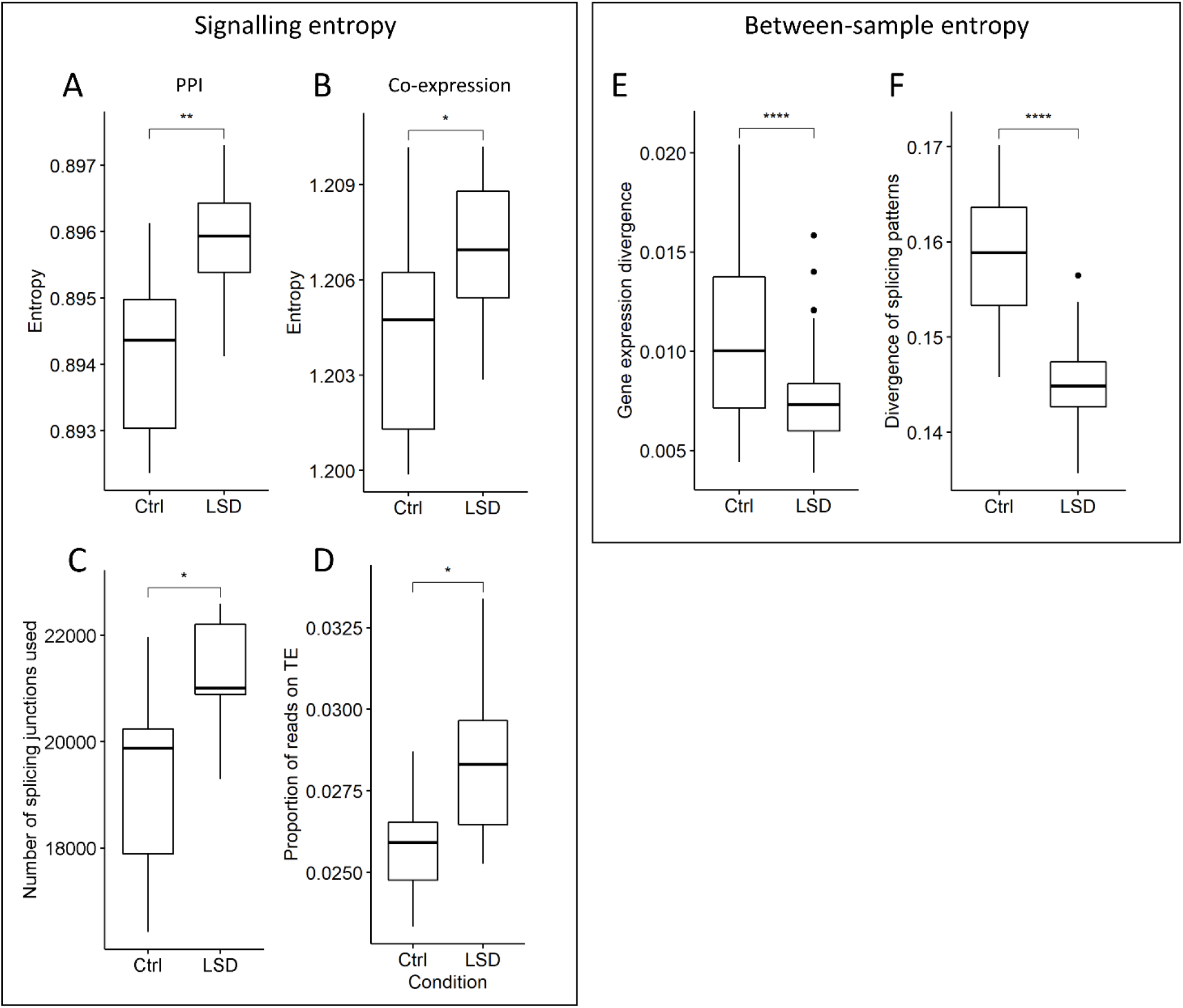
Signalling entropy increases in the LSD condition, while between-sample entropy decreases. A) Signalling entropy based on the PPI network; B) signalling entropy based on the co-expression network; C) signalling diversification quantified as the number of different splicing junction used in each sample or D) from the proportion of reads in each library mapping to transposable elements. Between-sample entropy measured from the correlation between samples using E) gene expression or F) splicing junction usage. Significance is obtained with the Wilcoxon rank sum test. * = 0.05, ** = 0.01, *** = 0.001, **** = 0.0001

### 2.3. Co-expression networks reorganize toward a less centralized topology

We next studied which nodes most strongly contribute to the overall signalling entropy increase, ranking the genes based on their average entropy change between the LSD and saline conditions.

Performing a GSEA analysis on the ranked gene list, RNA processing and splicing, chromosome organization, DNA repair, cell cycle, extracellular matrix, and chromosome organization resulted as significantly enriched amongst the genes with the highest increase in entropy. Of note, increased entropy in genes regulating RNA processing and splicing could explain the larger set of splicing junctions detected in the LSD treatment group. Interestingly, the genes with altered splicing patterns are enriched for genes regulated by the splicing factor Nova.

In line with an increase in entropy, the overall connectivity of the co-expression network decreases (Figure 4A), as previously shown for cancer networks (Schramm et al., 2010). Interestingly, not all modules rearrange their connections in the same way: despite most modules decreasing their co-expression in the LSD treated group, indicating a de-centralization of the network, three display increased intramodular connectivity, indicating a tighter regulation of corresponding functions (Figure 4B). For the same three modules, the nodes that increase co-expression connections tend to be the most central, while for the remaining modules we observe the opposite trend (Figure 4C,D). Accordingly, in most modules entropy increases more strongly for peripheral nodes, and only a few modules make an exception (Figure 5D). Of these, the “mesenchyme development” module shows coherent correlation sign between centrality and change in connectivity/entropy. This indicates that for most modules the nodes increasing signalling connections, reflecting on both entropy and intra-modular weighted degree, are the most peripheral in the module, while hubs tend to lose connections (Figure 5). Intriguingly, the modules showing strong opposite trends, and therefore increasing the compactness and centralization of the network, are enriched for mesenchyme development, regulation of immune system and extracellular matrix, categories that could reflect microenvironmental reorganization, and suggesting that different cell types might be affected by prolonged LSD treatment in different ways.

**Figure 4.**
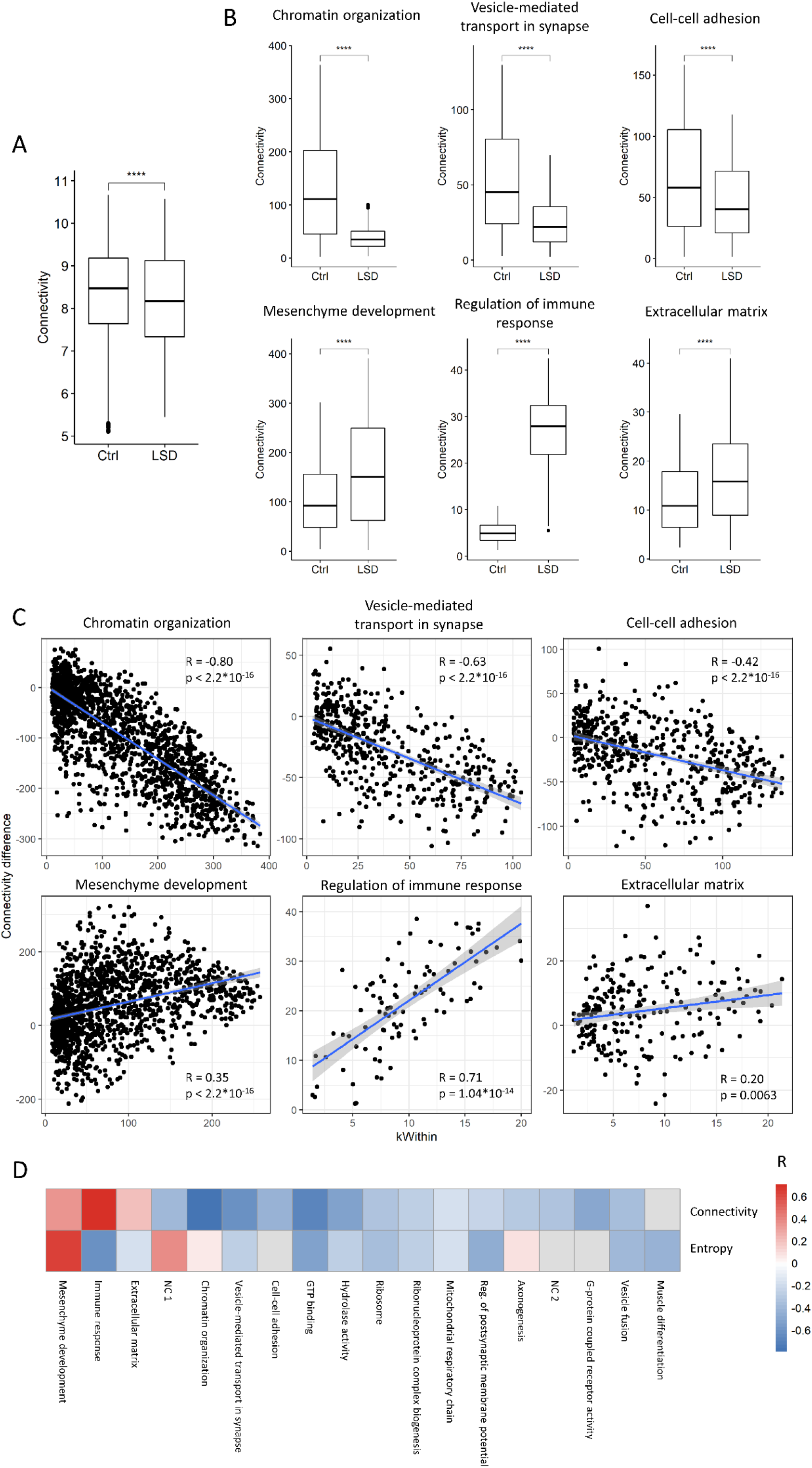
Different modules differently reorganize their structure upon LSD. A) The overall network connectivity decreases in the LSD condition; B) connectivity changes in the LSD-induced modules and in three additional modules showing opposite trend; C) Gene-wise relationship between change in connectivity and network centrality, measured as the intramodular connectivity (kWIthin). Together with each scatter plot, Pearson’s correlations and p-values are reported. D) Heatmap summarizing the correlation between node’s centrality and connectivity change (first row) or entropy change (second row) for each module. Red indicates positive and blue indicates negative correlation. Non-significant correlations are shown in grey.

**Figure 5.**
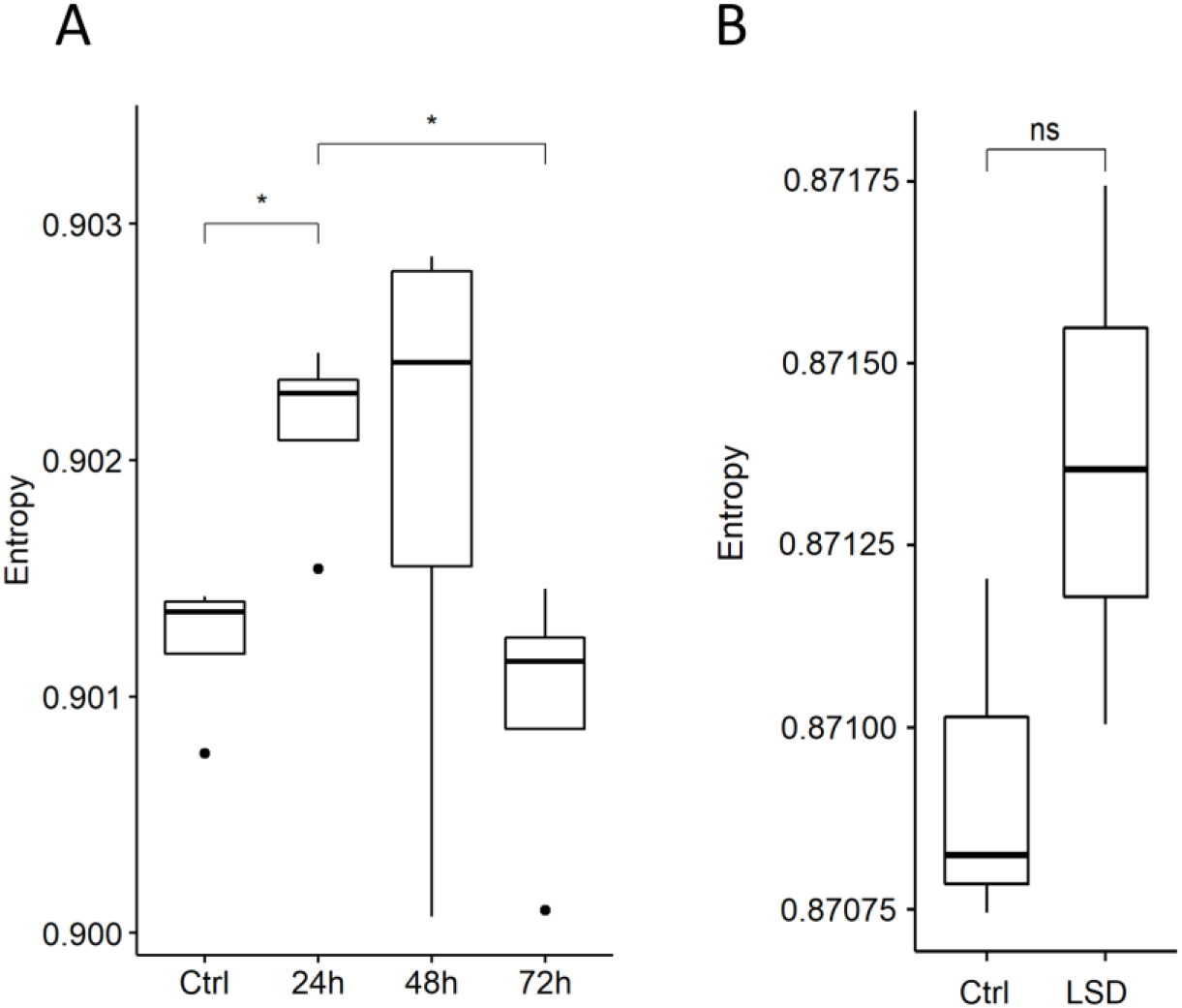
Signalling entropy transiently increase upon a single psychedelic’s administration. A) Signalling entropy of mice’s PFC neurons at different time points after a single DOI’s administration; B) Signalling entropy in rat’s PFC 90min after LSD treatment. Significance is obtained with the Wilcoxon rank sum test. * = 0.05, ns = not significant.

### 2.4. Persisting effects of repeated treatment resemble the transient effects of a single treatment

To explore the temporal dynamics of the transcriptional changes that we observe in chronically treated rats, we took advantage of two additional transcriptomic datasets, analysing PFCs from rat brains harvested 90 min after an acute administration of LSD (C. D. Nichols & Sanders-Bush, 2004), or neurons isolated from the PFC of mice after 24h, 48h or 72h of DOI administration (Revenga et al., 2021).

We envisioned at least two possible temporal dynamics: in the first model, psychedelics induce the well-established transcriptional changes in neuroplasticity genes without immediately affecting the organization of co-expression modules nor transcriptional complexity, exerting this effect only after repeated treatments; in a second model, the network changes that we detect are induced acutely, and potentially show different post-treatment dynamics in the case of single or multiple treatments.

Therefore, we tested whether our findings could be replicated in the two aforementioned single-dose datasets. Co-expression modules increasing in expression with chronic LSD treatment are strongly over-expressed also after a single treatment with DOI, and return to the baseline at 72h (Suppl. Fig. 2). Additionally, the potential transcriptional regulator of the “vesicle transport” module, Tcf4, shows the same trend upon DOI treatment (Suppl. Fig. 3A). Modules down-regulated upon chronic LSD treatment do not show a clear pattern, with only a slight trend toward down-regulation at 24h and 48h of the GTP-binding module (Suppl. Fig. 2). This suggests that, while the first three groups of genes are induced by acute treatment and require multiple administrations for sustained expression, the last three modules are repressed only upon chronic treatment. No significant differences could be detected in the microarray of LSD’s acute effects, possibly related to the small sample size (N=3), to differences in the technologies, or to the smaller number of genes analysed in the array with respect to genome-wide RNA-seq (Suppl. Fig. 3B, 4). Nevertheless, we cannot exclude the lack of differential expression being due to differences in the biological system.

Similarly, we tested the change in signalling entropy: in the LSD dataset, we observed a trend towards increased signalling entropy in the treated group, which nevertheless did not reach significance, likely due to the small sample size, while entropy is significantly increased by DOI at 24h after treatment, returning to baseline at 72h. Unfortunately, the small number of samples in the two additional datasets did not allow to have a reliable measure of between-sample entropy.

## 3. Discussion

Psychedelic drugs are gaining attention from the scientific community as potential new compounds for the treatment of psychiatric diseases such as depression and PTSD, but also as highways to explore the neurobiology of human consciousness. Also, the psychedelics’ induced alterations in perception and thought processes mimic symptoms of mental disorders such as schizophrenia, and hence they have been employed in animal models to study this disease. Neuroimaging studies have shown their effects on brain connectivity networks in humans, where they induce the reorganization of interacting areas leading to a diminished activity of the DMN and to an increased variability of the electromagnetic signal, quantified as an increase in the entropy.

The main molecular target has been identified as the serotonin 5-HT_2A_ receptor. Early studies pointed to an effect on neuroplasticity-related genes, but an extensive investigation on psychedelics’ induced gene expression changes is still lacking. Nevertheless, gene expression patterns and interactions strongly characterize the functioning of biological systems, implying that behavioural changes are often mediated by alterations in gene expression. The transcriptome, studied through microarray or RNA-sequencing, is the most accessible layer of gene expression, and RNA-seq is increasingly employed to explore the molecular mechanisms implicated in physiological, pathological and drug-induced processes.

Analysing RNA-seq data from prefrontal cortices of rats chronically treated LSD three months after drug discontinuation (e.g. no drug or recent drug on-board), we observe long-lasting changes in gene expression, particularly in neuroplasticity and neurotransmission genes, in line with previous reports, but also in circadian rhythm and, importantly, in epigenetic modifiers. Indeed, Tet1, involved in the erasure of DNA methylation (Wu & Zhang, 2017), increases its expression upon LSD, and the “histone modification” GO category showed strong enrichment for up-regulated genes. This indicates that repeated psychedelics’ administration affects the epigenetic machinery, thus proposing a mechanism for their long-term effects. Interestingly, Tet1 is recruited by Egr1 (Sun et al., 2019), one of the classic immediate-early genes which expression is induced by psychedelics (González-Maeso et al., 2003). Of note, it has been recently shown that a single dose of DOI is able to alter the PFC epigenetic status in mice up to 7 days after administration (Revenga et al., 2021), suggesting that a single exposure is sufficient to induce epigenetic reorganization.

Applying gene co-expression network analysis, we confirm these observations identifying modules of tightly connected LSD-induced genes enriched for chromatin modification, vesicle mediated transport in synapse and cell-cell adhesion. On the other hand, down-regulated modules are enriched for GTP-binding, hydrolase activity and ribosome. From the topology of the co-expression network, we propose Tcf4 as a potential regulator of the up-regulated “Vesicle transport” module. Notably, Tcf4 regulates neuritogenesis and neuronal migration, and its loss decreases spine density in the cortex and in the hippocampus (Crux et al., 2018).

Importantly, we describe an overall increase in transcriptional/signalling entropy, potentially reflecting an overall increase in available transcriptional states. Intriguingly, transcriptional entropy mirrors, at the molecular level, the increased brain entropy reported through neuroimaging studies in human.

High transcriptional entropy is typical of stem cells, and decreases with differentiation, paralleling cells’ developmental potential. Nevertheless, transcriptional entropy has also been associated with aging, reflecting the disruption of a well-organized and functional system.

Using multiple metrics, we show that LSD-induced transcriptional entropy is more reflective of a plastic stem-like state than an aged state, suggesting the induction of a potentiality-expansion process with organized and reproducible features. Moreover, we imply additional players in transcriptional diversification, since LSD-treated rats display re-activation of transposable elements and increased alternative splicing sites’ usage. In particular, the most reliably alternatively spliced genes are enriched for targets of the Nova splicing factor, known to control the splicing of synaptic proteins (Ule et al., 2005).

Transposable elements (TEs) are a class of repeated DNA sequences with the ability to mobilize and change locations in the genome (Ahmadi et al., 2020). Despite being mostly inactive in somatic cells, efficient transposition was detected in neural progenitor cells and mature neurons (MacIa et al., 2017). They have been reported to be both beneficial and pathological to the organism (Biémont, 2010), and associated with both neurodegenerative and psychiatric disease and plasticity. For example, in first episode schizophrenia, hypomethylation of HERV-K locus was reported (Forner et al., 2019), and L1 insertions were found significantly elevated in post-mortem dorsolateral prefrontal cortex of patients with schizophrenia (Doyle et al., 2017). Nevertheless, they have been proposed to have a fundamental role in promoting evolution and also increasing cells’ variability through the generation of somatic mosaicism (Paquola et al., 2017), creating a greater potential for the adaptation of genetic networks.

Disentangling individual modules’ topological reorganization, we distinguish two opposite processes: 1) most modules decrease their connectivity paralleling the increased entropy, and redistribute connections towards module’s periphery, hence reducing its centralization (Figure 6A); 2) a few modules increase their overall connectivity and centralization. Interestingly, the modules increasing connectivity are enriched for GO categories indicative of the involvement of microenvironment: mesenchyme development, regulation of immune response and extracellular matrix. This suggests a potential involvement of non-neuronal cells in long-term LSD effects. In particular, Mesenchymal stem cells (MSCs) are stem cells found in many adult tissues, including brain, (Appaix, 2014) and can differentiate into neurons (Pavlova et al., 2012; Zeng et al., 2015; Zhao et al., 2015) and glial cells (George et al., 2019) to replace damaged tissues (Dimarino et al., 2013), and thus promoting neuroprotection, regeneration and repair (Qu et al., 2008; Thomi et al., 2019). They have immunomodulatory properties, mitigating the inflammation related to stroke or neurological diseases (Feng et al., 2020; Liu et al., 2019; Salari et al., 2020). Interestingly, both mesenchyme- and immune-related modules display similar topological restructuring.

**Figure 6.**
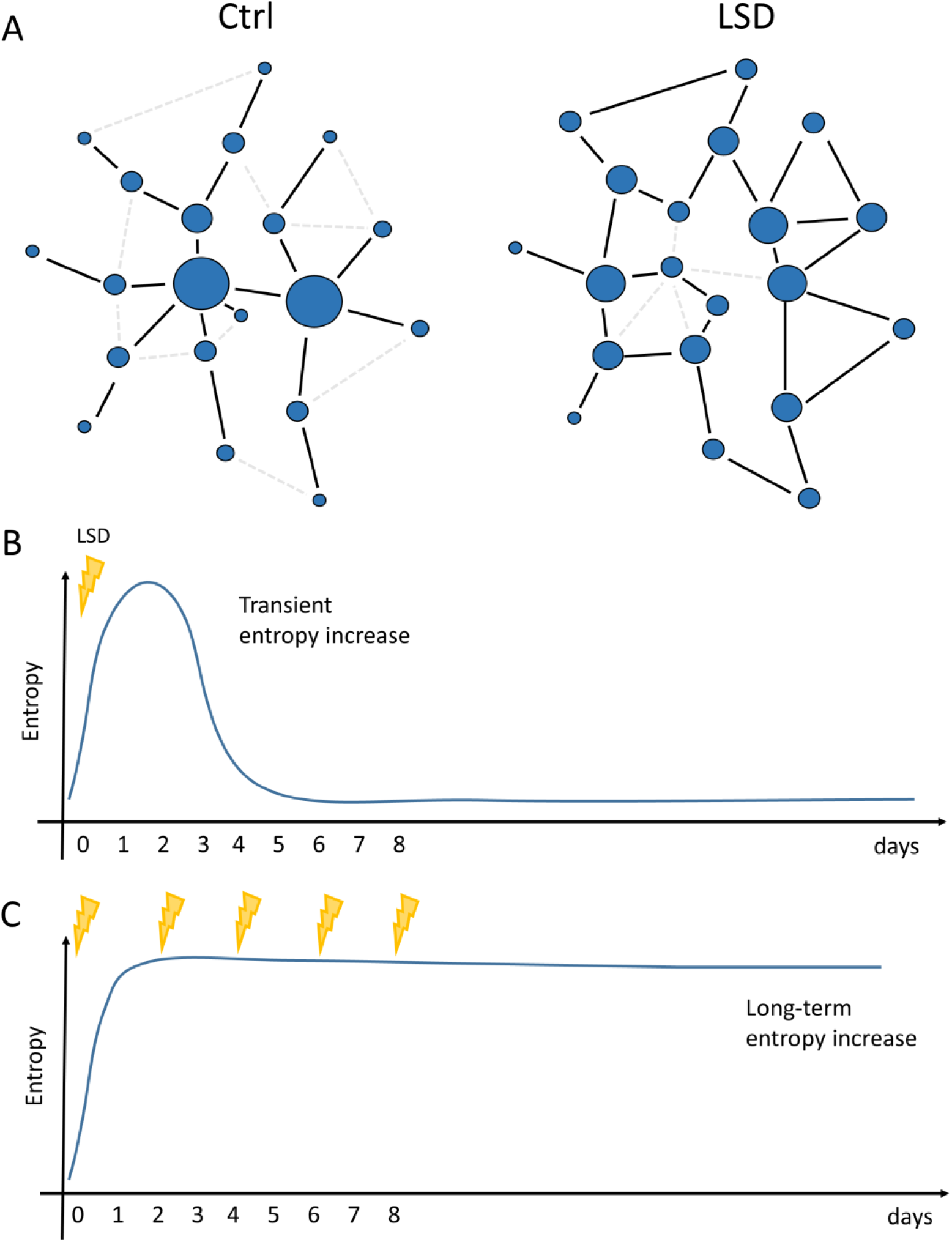
Schematic representations of models of LSD-induced changes in network organization. A) Most gene co-expression modules reorganize toward a less centralized structure, with central hubs losing connections and peripheral nodes acquiring connections. Overall, the number of available paths across the network increases, resulting in increased signalling entropy. In the proposed model, a single dose of psychedelic drug induces a transient increase in signalling entropy that returns to baseline after 72h (B), while repeated administrations induce a sustained entropy increase even after discontinuation of the drug (C).

In line with our observations suggesting non-neuronal populations’ involvement, in particular of the immune system, psychedelics have been shown to exert anti-inflammatory effects, which led to their proposal as treatments for neurodegenerative diseases such as Alzheimer disease (Family et al., 2020; Flanagan & Nichols, 2018; Vann Jones & O’Kelly, 2020).

Finally, we explored the temporal dynamics of co-expression modules and signalling entropy by comparing our results with other available datasets. We acknowledge that none of these conditions is ideal for our aim, since the LSD dataset has been generated with a different platform (Affymetrix U34A), possibly generating technical batch effects, and comprises only three pooled independent replicates of two animals each, limiting the statistical power, while the second utilizes a different psychedelic drug, DOI, with non-perfectly overlapping effects with LSD (González-Maeso et al., 2003). Another limitation is that the LSD set used rats, and the DOI set mice. Nevertheless, both LSD-induced modules and signalling entropy display similar trends in PFC neurons upon DOI treatment, increasing at 24h and returning to background after 72h. Hence, we propose a model of the transcriptional response to psychedelics where a single dose triggers a transient reorganization of the gene networks that is sustained at long-term only after several administrations (Figure 6B).

We hypothesise that the short-term reorganization and entropy increase induced by LSD allows for the formation of new synaptic connections and hence novel neuronal networks that can be maintained after the treatment, resulting in the long-lasting beneficial effects that psychedelics have proven to exert in clinical settings. Frequent and repeated administrations, however, may result in a prolonged increase in the entropy and decentralization of co-expression networks that could be reflective of the psychotic-like state observed in the chronically treated rats.

Signalling entropy increase could be interpreted as increased cell diversity within the same subject (Nijman, 2020), but could also reflect higher individual cells’ potential. With the available data, it is impossible distinguishing between these two mechanisms, for which single cell RNA-seq experiments would be necessary (Papalexi & Satija, 2018). The indirect activation by LSD of several neuronal cell types has previously been shown (Martin & Nichols, 2016), but how each cell population responds to this class of compounds and how cell heterogeneity is affected remains largely unknown.

Additionally, comparing experiments with different administration schedules showed differences in the persistence of transcriptional rewiring, and additional time-course experiments with longer time frames will be needed to elucidate the dynamics of gene expression response.

In conclusion, analysing transcriptomic data of LSD treated rats, we suggest some of the possible molecular mechanisms potentially underlying psychedelics-induced increase in brain entropy observed at higher psychedelic levels, identify epigenetic alterations that could explain long-term effects, and imply alternative splicing, transposable elements’ activity and the involvement of additional cell components such as the mesenchyme in the mechanism of action of psychedelics.

## 4. Methods

### 4.1. Data collection and pre-processing

#### RNA-seq data of rats chronically treated with LSD

Data collection is described in (Martin et al., 2014). Reads were trimmed and quality checked with TrimGalore (https://www.bioinformatics.babraham.ac.uk/projects/trim_galore/), aligned to the Rattus norvegicus genome (rn6) with Hisat2 (Kim et al., 2019), and counts obtained with HTSeq-count (Anders et al., 2015). Resulting processed data were RPM normalized and only genes with at least 5 reads in at least 10 samples were retained for further analyses.

#### Microarray data of rats after 90min of LSD treatment

Experimental setting and data collection are described in C. D. Nichols & Sanders-Bush, 2004. Raw data were normalized with the rma function from the Affy package (Gautier et al., 2004). Probes were mapped to gene symbols with the GPL85 annotation from Gene Expression Omnibus.

#### RNA-seq of mice’s cortical neurons upon treatment with DOI

Counts were downloaded from Gene Expression Omnibus (GSE161626). Technical replicates were averaged and then normalized and log transformed similarly to rats’ PFC data.

Splicing junctions were quantified with SGSeq (Goldstein et al., 2016). Transposable elements were quantified with TEtranscripts (Jin et al., 2015), after aligning the reads with Hisat2 allowing reporting up to 1000 mapping sites per read.

### 4.2. Subsampling

Subsampling was performed by randomly selecting 10 million reads from the original fastq files, and repeating the pre-processing, as described above.

### 4.3. Differential expression and functional enrichment

Differential gene expression was performed with DESeq2 (Love et al., 2014) on count data. Gene Ontology enrichment was calculated with the enrichGO function from the clusterProfiler package (Yu et al., 2012), using “Biological Process” or “Molecular Function” GO categories and default parameters. For modules’ enrichment, all genes belonging to a module different from the “grey” (unconnected) module were used as background.

Transcription Factors (TFs) were defined based on the Gene Ontology category GO0003700, downloaded from Biomart (https://www.ensembl.org/biomart/).

Targets of each TF were obtained from the ChEA database, and downloaded from https://maayanlab.cloud/Enrichr/#stats (ChEA_2016).

An exploratory enrichment analysis of differentially spliced genes has been performed with Enrichr (https://maayanlab.cloud/Enrichr/). The Wikipathway gene lists has been downloaded from https://maayanlab.cloud/Enrichr/#stats (Wikipathway_2021_Human) and used for testing the enrichment of significantly differentially spliced genes.

The GSEA analysis of genes with the highest entropy change was performed with the fgsea (Korotkevich et al., 2021) and msigdbr packages (https://CRAN.R-project.org/package=msigdbr), using the category “C5” (comprising GO categories).

### 4.4. Co-expression networks

The co-expression network was obtained with the blockwiseModules function from the WGCNA package (Langfelder & Horvath, 2008), setting the parameters networkType and TOMtype to “signed”, and beta=12.

The module eigengene (ME) was calculated with the function moduleEigengenes, and weighted degree within each module (kWithin) was calculated with the function intramodularConnectivity.fromExpr, setting networkType=“signed” and power=12.

Differential activity of modules was tested comparing ME between treatment groups with the Wilcoxon test, and then correcting for multiple testing and obtaining false discovery rates for each module.

Connectivity represents genes’ weighted degree, with weights based on the adjacency matrix, either across the whole network or considering only a module’s genes. The connectivity in a specific condition was obtained by calculating the adjacency using only samples belonging to that group

Module eigengene in different datasets were obtained as the projection of samples on the first PC of a PCA (function prcomp) performed on rats’ PFC data using only the genes of the chosen module.

### 4.5. Signalling entropy

Signalling entropy was calculated with the SCENT R package (Teschendorff & Enver, 2017), either using the provided PPI network or the co-expression network built in this work. Input data (RNA-seq) were log transformed with an offset of 1.1.

### 4.6. Between-sample entropy

Between-sample entropy was obtained as 1-correlation, with the correlation obtained from pairwise Pearson’s correlation between samples using gene expression profiles or splicing junction usage profiles.

### 4.7. Gene mapping

ID mapping and orthologs identification was performed with the biomaRt R package (Durinck et al., 2009). In case of multiple IDs mapping to the same symbol, the ID with the highest average expression across all dataset’s values was chosen.

### 4.8. Statistical analyses

All statistical analyses were performed with R 4.0.4 (R Core Team, 2018).

Packages used for plotting are R base graphics, ggplot2 (Wickham, 2017) and ggsignif (https://CRAN.R-project.org/package=ggsignif), and pheatmap (https://CRAN.R-project.org/package=pheatmap).

Groups’ means were compared with the Wilcoxon rank sum test, while correlations’ statistical significance was calculated with the cor.test function. Only tests passing the threshold FDR<0.05 (false discovery rate) are shown.

## Supplementary material

**Supplementary Figure 1.** Changes in signalling entropy (A,B) and number of splicing junctions used (C) do not depend on sequencing depth. Results shown have been obtained through subsampling of original sequencing data to the same depth of 10 million reads.

**Supplementary Figure 2.** Modules regulated upon LSD treatment in the DOI treatment dataset. Significance is obtained with the Wilcoxon rank sum test. * = 0.05, ** = 0.01, *** = 0.001, **** = 0.0001

**Supplementary Figure 3.** Tcf4 regulation in the DOI (A) and acute LSD treatment (B) datasets. Significance is obtained with the Wilcoxon rank sum test. * = 0.05, ** = 0.01, *** = 0.001, **** = 0.0001

**Supplementary Figure 4.** Modules regulated in the acute LSD treatment dataset. Significance is obtained with the Wilcoxon rank sum test. * = 0.05, ** = 0.01, *** = 0.001, **** = 0.0001 **Supplementary Table 1.** Differentially expressed genes upon treatment with LSD

**Supplementary Table 2.** Gene Ontology categories enrichment for up-regulated genes after treatment with LSD.

**Supplementary Table 3.** Gene Ontology categories enrichment for down-regulated genes after treatment with LSD.

**Supplementary Table 4.** List of genes belonging to each co-expression module.

**Supplementary Table 5.** Gene Ontology Biological Processes enrichment for co-expression modules’ genes.

**Supplementary Table 6.** Gene Ontology Molecular Function enrichment for co-expression modules’ genes.

