## Supplementary figures and images for "LSD induces increased signalling entropy in rats’ prefrontal cortex"

### Supplementary Figure 1

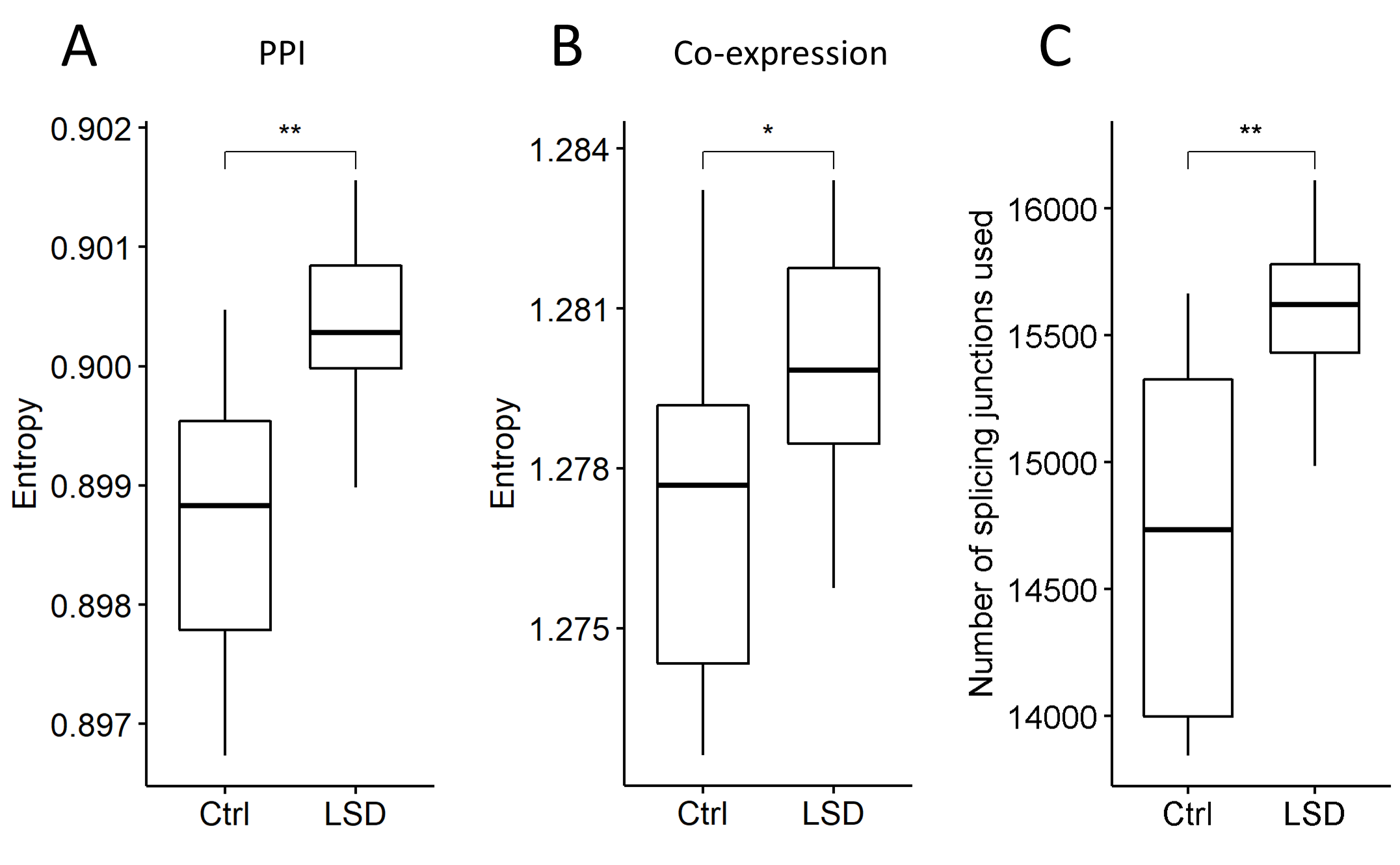

### Supplementary Figure 2

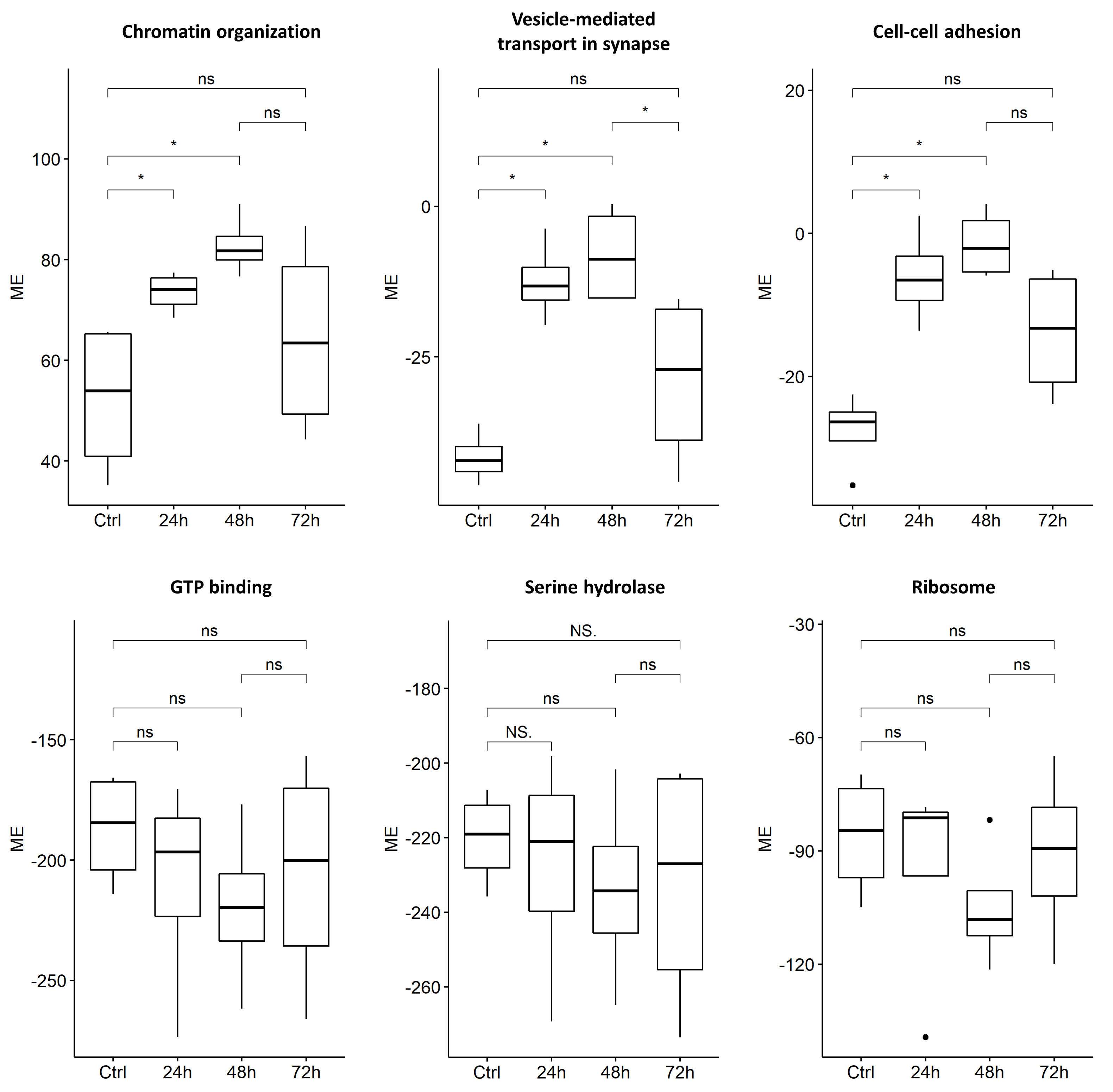

### Supplementary Figure 3

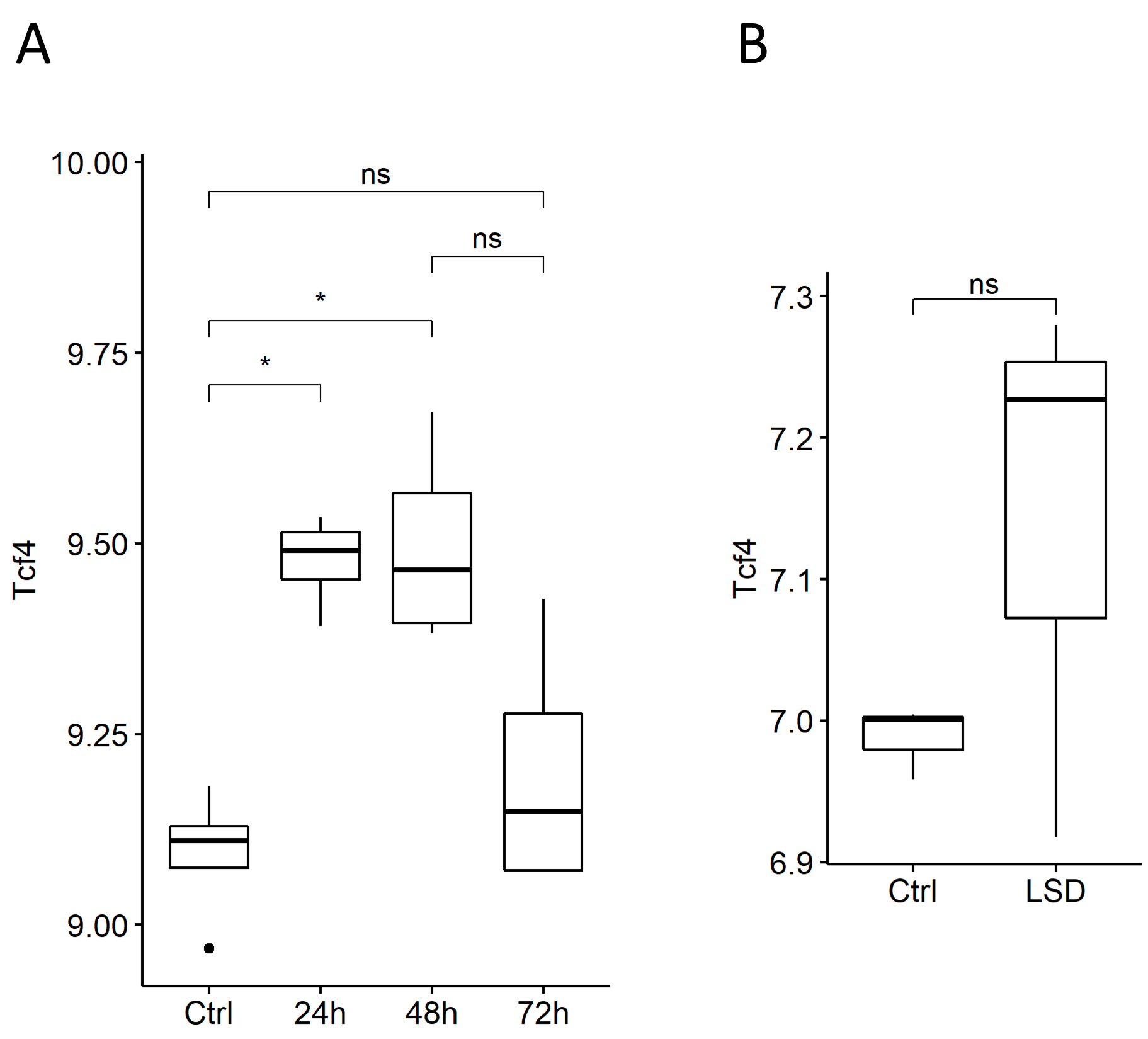

### Supplementary Figure 4

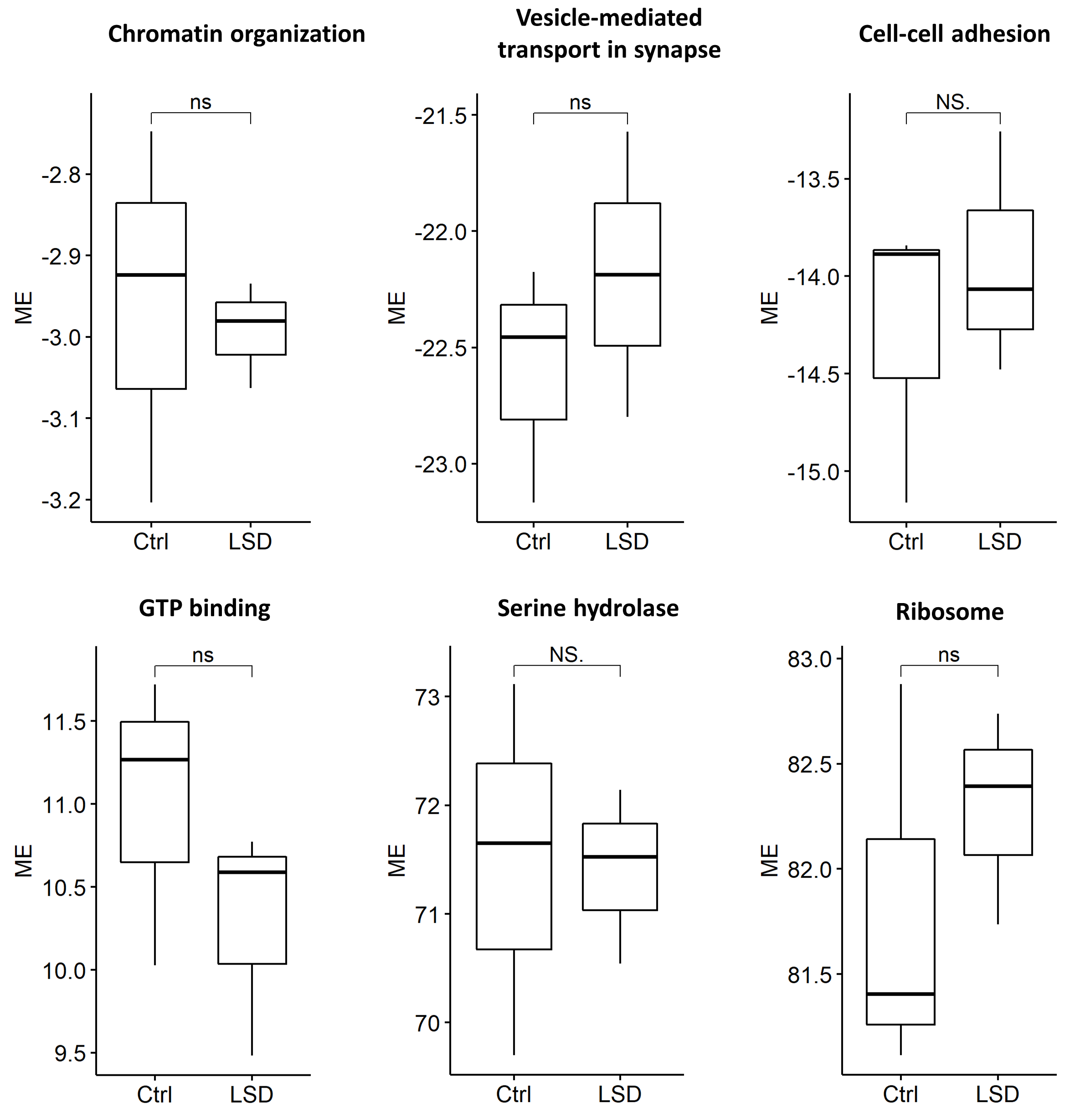
